# Diterpenoid phytoalexins shape rice root microbiomes and their associations with root parasitic nematodes

**DOI:** 10.1101/2024.09.30.615782

**Authors:** Enoch Narh Kudjordjie, Willem Desmedt, Tina Kyndt, Mogens Nicolaisen, Reuben J. Peters, Mette Vestergård

## Abstract

Rice synthesizes diterpenoid phytoalexins (DPs) are known to operate in defense against foliar microbial pathogens and root-knot nematode *Meloidogyne graminicola*. Here, we examined the role of DPs in shaping rice⍰associated root microbiomes in nematode-infested rice paddy soils. Further, we assessed how DPs affect interactions between the root microbiomes and nematode communities, particularly rice root-knot nematodes from the *Meloidogyne* genus. We used 16S and ITS2 rRNA gene amplicon analysis to characterize the rice root-and rhizosphere-associated microbiomes of DP knock-out mutants and their wild-type parental line, at an early (17 days) and late (28 days) stage of plant development in field soil. Disruption of DP synthesis resulted in distinct composition and structure of microbial communities both relative to the parental/wild-type line but also between individual mutants, indicating specificity in DP-microbe interactions. Moreover, nematode-suppressive microbial taxa, including *Streptomyces*, *Stenotrophomonas*, *Enterobacter*, *Massilia*, and *Acidibacter*, negatively correlated with *Meloidogyne*. Comparative analysis revealed differential enrichment of microbial taxa in the roots of rice diterpenoid phytoalexin (DP) knock-out mutants versus wild-type, suggesting that DPs modulate specific taxa in the rice root microbiome. These findings indicate DPs role in plant-microbiome assembly and nematode interactions, further underscoring the potential of leveraging phytoalexins for sustainable management of crop diseases.

## Introduction

Plants constantly must defend themselves against biotic and abiotic stressors. To protect themselves against microbial pathogens or parasitic nematodes, plants as sessile organisms have adapted constitutive and inducible chemical repertoires that are part of the conserved immune system (Chisholm *et al*., 2006). Inducible defense responses are mediated in part by stress-inducible secondary metabolites collectively called phytoalexins. Phytoalexins are a structurally diverse class of low weight molecules produced by the host in response to biotic and abiotic stressors (VanEtten *et al*., 1994). Several classes of phytoalexins, including terpenoids (diterpenoids, triterpenoids, sesquiterpenoids), phenylpropanoids and benzoxazinoids, have been identified in different plant families, with profound protective roles against microbial pathogens, nematodes and abiotic stresses (Dixon, 2001; Lu *et al*., 2018; Sikder *et al*., 2021; Desmedt *et al*., 2022; Kaur *et al*., 2022). The ability of phytoalexins to defend the host against plant pathogens is being investigated as a potential target for the sustainable control of plant diseases (Jeandet *et al*., 2017).

Diterpenoids are among the major classes of phytoalexins produced in several monocots including rice (*Oryza sativa*) (Yamane, 2013; Murphy and Zerbe, 2020), in which the predominant phytoalexins include the diterpenoid momilactones, phytocassanes, and abietoryzins (Schmelz *et al*., 2014; Kariya *et al*., 2020, 2023). Biosynthesis of the rice diterpenoid phytoalexins (DPs) proceeds via either *ent*- or *syn*-copalyl diphosphate (CDP) intermediates produced from the general diterpenoid precursor geranylgeranyl diphosphate (GGDP) (Yamane, 2013; Schmelz *et al*., 2014). GGDP undergoes cyclization to form either *ent*-CDP or *syn*-CDP, catalyzed by CDP synthases (CPSs), with *syn*-CDP exclusively produced by OsCPS4, and *ent*-CDP produced by OsCPS1 and OsCPS2 (Otomo *et al*., 2004; Prisic *et al*., 2004; Xu *et al*., 2004). OsCPS1 is specifically required for gibberellin phytohormone biosynthesis (Sakamoto *et al*., 2004), while the expression of OsCPS2 is inducible and associated with DP biosynthesis (Toyomasu *et al*., 2008). Subsequently, rice kaurene synthase-like (OsKSL) enzymes convert *syn*-CDP or *ent*-CDP into olefin precursors of the various groups of DPs (Yamane, 2013; Schmelz *et al*., 2014).

Rice DPs are produced in different plant organs and accumulate during stress conditions such as exposure to UV or pathogen attack (Yamane, 2013; Schmelz *et al*., 2014). Rice roots exude DPs such as momilactones and phytocassanes into the rhizosphere (Toyomasu *et al*., 2008). These DPs have a broad spectrum of biocidal activities against microbial pathogens such as *Magnaporthe oryzae* and *Xanthomonas oryzae* (Toyomasu *et al*., 2008). In addition, terpenoid phytoalexins in other plants have been shown to modulate the assembly of their associated microbiomes (Huang *et al*., 2019; Murphy *et al*., 2021). For example, Li et al., (Li *et al*., 2023) <u>reported</u> differential inhibitory effects of distinct switchgrass DPs on fungal taxa in their rhizosphere, and Murphy et al. (Murphy *et al*., 2021) recently showed that maize-secreted DPs affect bacterial communities in the rhizosphere.

Our previous work with rice DP-deficient mutant lines grown in field soil demonstrated that these secondary metabolites protect rice plants against the root-knot parasitic nematode *Meloidogyne graminicola*, likely through a combination of direct nematostatic activity and alterations in root-associated nematode communities that include the attraction of predatory nematodes believed to feed on phytoparasitic nematodes (Desmedt *et al*., 2022). In addition to the direct effects of DPs on root nematodes, these secondary metabolites may modulate the root microbiome in ways that alter susceptibility to nematode pests. Root microbiomes assembled by other plants during pathogen or pest attack have been shown to encompass beneficial microbes that are able to suppress plant-parasitic nematodes through the secretion of secondary metabolites and lytic enzymes (Chinheya *et al*., 2017; Poveda *et al*., 2020). Examples include rhizobacteria such as *Bacillus* spp., *Rhizobium* spp., *Burkholderia spp*., *Pseudomonas* spp., and the fungi *Trichoderma* spp. and *Clonostachys rosea* (Chinheya *et al*., 2017; Iqbal *et al*., 2018; Poveda *et al*., 2020). Therefore, in this study we sought to examine the effect of DPs on rice-associated microbiomes assembled in *M. graminicola* infested field soil.

In particular, we hypothesize that DPs not only increase rice resistance towards *M. graminicola* by directly affecting nematodes but also acts through effects on the host-root associated microbiomes. We therefore determined the effects of rice DP disruption mutants on the composition of the root microbiome and examined the correlation between the microbiota and nematode communities, specifically focusing on *M. graminicola*.

## Materials and methods

### Plant material

We used the rice wild-type (WT) *Oryza sativa*L. subsp*. japonica* cv. Kitaake and derived DP-deficient CRISPR-Cas9 knock-out mutants disrupted in *OsCPS2* (*cps2*), *OsCPS4* (*cps4*) and the double mutant *cps2/4*. In *cps2*, the accumulation of *ent*-CPP derived DPs is significantly reduced, and in *cps4*, the production of *syn*-CPP derived DPs is blocked. The DP profiles of this parental/wild-type and these mutants, as well as their lack of effect on plant growth and development, have been previously described (Zhang *et al*., 2021).

### Experimental design and sampling

Experimental design, growing of plants, sample collection and DNA extraction from rice lines were performed as previously described (Desmedt *et al*., 2022). Briefly, rice lines were grown in pots filled with 250 ml of *M. graminicola* infested field soil with a long history of rice cultivation, collected on August 26^th^, 2021 in the commune of Garlasco (Pavia province, Lombardy, Italy). Each pot was planted with one rice seedling (5 days after placing on wet tissue paper), representing a biological replicate. Seedlings were placed in a greenhouse at 26°C and 12 h of artificial light per day for the entire duration of the experiment. During the first 24 hours after transplantation to the pots, the seedlings were covered with a transparent plastic sheet. The pots were irrigated with 70 ml of distilled water at transplantation and then with 15 ml distilled water daily. We fertilized the pots twice, at 1 and 14 days post transplantation (dpt), with 15 ml of a fertilizer solution containing 2 g l^-1^ ferrous sulfate heptahydrate and 1 g l^-1^ ammonium sulfate. Seven days after transplantation, we inoculated the soils with an additional 120 *M. graminicola* infective stage J2s, to ensure a high probability of root infestation.

We sampled bulk soil, rhizosphere soil and roots at two times points, at 17 d and 28 d after transplantation (dpt), as described previously (Desmedt *et al*., 2022). We carefully uprooted five seedlings from each line and gently shook the roots to remove nonadherent soil after which adhering rhizosphere soil was gently scraped off and flash-frozen in liquid nitrogen. We then washed the root systems thoroughly with distilled water and froze the root separately. The collected samples were kept frozen, lyophilized for 48 h, and prepared for DNA extraction by grinding with sterile steel beads at 3×1500 strokes per min in a GenoGrinder (Ramcon, Birkerød, Denmark). We extracted DNA from soil samples using a Qiagen PowerLyzer Soil kit, and from root samples using a Qiagen DNeasy Plant Pro kit. DNA concentrations were checked using a Qubit fluorometer (Thermo-Fisher Scientific, Waltham, MA, USA), prior to downstream processing.

### Amplicon library preparation

To characterize microbial communities, we amplified and sequenced bacterial 16S and fungal ITS regions by following a previously described procedure (Kudjordjie *et al*., 2019). Briefly, bacterial amplicon libraries were constructed by amplifying 16S rRNA V5-V7 with 799F/1193R primer pairs (Chelius and Triplett, 2001; Bodenhausen *et al*., 2013; Beckers *et al*., 2016). Bacterial amplification was performed in a 25 μl reaction mix consisting of 12.5 μl Invitrogen Platinum SuperFi PCR master mix (Thermo Fisher Scientific, Waltham, Massachusetts), 2 μl of each primer (10 μM stock), and 6.5 μl nuclease free water, and 2 μl of the template DNA. PCR amplification was performed in a GeneAmp PCR System 9700 thermal cycler (Thermo Fisher Scientific) at the following conditions: 94 °C for 5 min; 25 cycles at 94 °C for 30 s, 55 °C for 30 s and 72 °C for 30 s; and a final extension step at 72 °C for 10 min. The fungal library was prepared using the fITS7 and ITS4 fungal primer pair that amplify the internal transcribed spacer 2 (ITS2) region of the fungal rRNA gene (Ihrmark *et al*., 2012). Amplification was done in a reaction mixture of 25 μl consisting of 1× PCR reaction buffer, 1.5 mM MgCl2, 0.2 mM dNTPs, 1 μM of each primer, 1 U of GoTaq Flexi polymerase (Promega Corporation, Madison, USA), and 1 μl of DNA template. Temperature cycling conditions for fungal PCR were as described above, except for an annealing temperature at 57 °C (Ihrmark *et al*., 2012). We used dual indexing in combination with internal barcodes to pool samples. The bacterial and fungal forward primers were tagged with varying bases of multiplex identifiers (MID). For indexing, primers including indexing tags were used in a PCR for 10 cycles, with the thermal cycler programs as described above.

The index combinations fungal primer sequences are described in Kudjordjie et al. (Kudjordjie *et al*., 2019), while bacterial primer sequences including MID are presented in **Supplementary Table S1**. After PCR, amplicon sizes of both bacteria and fungi libraries were confirmed on 1.5% agarose gel, and bands of the expected sizes were excised and extracted using QIAquick Gel Extraction Kit (Qiagen). The libraries were sequenced on an Illumina MiSeq sequencer with PE300 at Eurofins MWG (Ebersberg, Germany).

### Sequence data and statistical analysis

Bacterial and fungal sequence reads were analyzed as described earlier (Kudjordjie *et al*., 2019). Briefly, paired-end reads were demultiplexed for internal barcodes, using Mr_Demuxy using command pe_demuxer.py (https://pypi.org/project/Mr_Demuxy/). Following this, the paired-end reads were assembled and joined using vsearch v.2.6 (Rognes *et al*., 2016). Primers were removed using cutapdapt (Martin, 2011), followed by dereplication, chimera removal, and operational taxonomic unit (OTU) clustering using vsearch v.2.6 (Rognes *et al*., 2016). Fungal ITS reads were extracted with ITSx extractor version 1.0.6 (Bengtsson-Palme *et al*., 2013) before clustering. Taxonomic assignments of bacterial and fungal OTUs were performed using the SILVA 132 (Quast *et al*., 2013) and the UNITE (v7.2) (Abarenkov *et al*., 2010)) databases, respectively, with assign_taxonomy.py in QIIME (v1.9)(Caporaso *et al*., 2010). The unassigned OTUs at kingdom level or OTUs assigned as chloroplast or mitochondrial sequences were removed.

Microbial analysis and visualizations were performed in R v4.0.5 (R Core Team, 2022), using phyloseq (v1.34.0.) (McMurdie *et al*., 2013), vegan (v2.5.7) (Oksanen *et al*., 2020), and ggplot2 (v3.3.2) (Wickham, 2009) packages. Sequences with less than 2000 and 1000 reads were removed from the bacterial and fungal data sets, respectively. For alpha diversity, OTU tables were rarified 100 times at a depth of 2000 reads for bacteria and 1000 reads for fungi and the mean of the diversity estimates of 100 trials was used to estimate observed species richness and Shannon diversity. Significant differences between alpha diversities were calculated using Kruskal-Wallis rank sum test. Before beta diversity analysis, OTU tables were transformed to relative abundances. The Bray–Curtis dissimilarity matrix was calculated to construct both unconstrained principal coordinates analysis (PCoA) and constrained principal coordinate (CAP) visualizations. CAP analysis was constrained by the host genotype using the function “ordinate” in the phyloseq package. Permutational analysis of variance (PERMANOVA) was computed on both bacterial and fungal datasets to quantify differences between experimental factors by using the adonis2 function from the “vegan” package (Oksanen *et al*., 2020). The analysis was performed using subset datasets of compartment (root, rhizosphere) and time points 17 and 28 days after transplantation (dpt). Further pairwise comparisons between WT and individual mutants were performed using the pairwise.adonis2 function (https://github.com/pmartinezarbizu/pairwiseAdonis/blob/master/pairwiseAdonis/R/pairwise.adonis2.R). Relative abundance of *M. graminicola* in rice mutants and WT were also computed to support nematode colonization of rice plants.

Microbial co-occurrence analysis was performed using a previously described procedure. Briefly, bacterial, fungal and nematode datasets (Desmedt *et al*., 2022) from 17 and 28 dpt were pooled, subjected to trimmed mean of M values (TMM) procedure using EdgeR (Robinson *et al*., 2010) in R. TMM-normalized data were used to calculate Spearman rank correlations between microbial OTUs using the corr function. We used OTUs that were present in at least 20% of the samples with r > 0.4 for positive correlations and r <−0.4 for negative correlations, and p-values < 0.01. The correlations were visualized in heatmaps.

Differential abundance analysis between WT and individual mutants was performed using ANCOM-BC (Lin and Peddada, 2020). ANCOM-BC utilizes linear regression and controlled FDR to identify differentially abundant taxa between rice-mutants and their parental wildtype.

## Results

To establish whether and how DPs modulate root-associated microbiomes in rice, we compared the bacterial and fungal communities of the parental/wild-type cultivar Kitaake and derived DP disrupted mutants. Bacterial 16S and fungal ITS regions were amplified and sequenced; the number of reads obtained per sample are summarized in **Supplementary Table S2** and the read distribution and rarefaction curves are given in **Supplementary Fig. S1**. The rarefaction curves were reaching an asymptote, reflecting satisfactory representation of microorganisms in the samples. Because microbial data was obtained from *M. graminicola* infested samples, we correlated the microbial data with the nematode sequence reads obtained from our previous study, which confirmed the role of DPs in rice defense against root-knot nematodes (Desmedt *et al*., 2022). Nematode sequence reads assigned to the genus *Meloidogyne* accounted for > 90% of reads (WT: 90%, cps2: 96%, cps4: 96%, cps2/4: 93%; P= 0.61), with strongest enrichment in roots at 17 dpt than 28 dpt. However, a significant difference was observed at the latter time point between wild-type and the *cps* mutants (Desmedt *et al*., 2022).

### Diterpenoids shape root and rhizosphere microbiomes

We found marked differences in the abundances of bacterial phyla and fungal classes in the roots of DP disruption mutants compared to WT (**Fig. 1A,B**). At 17 dpt, the relative abundances of bacterial phyla Proteobacteria were significantly higher in the rhizosphere of *cps4* and *cps2/4* compared with WT. Similarly, Firmicutes was higher in *cps4* than WT. In roots, the abundance of Actinobacteria was higher in the mutants than in WT at 17 dpt. Actinobacteria was highly enriched in *cps2* and *cps2/4*, while Firmicutes and Proteobacteria significantly decreased in *cps4* compared with WT. Also, Proteobacteria was lower in *cps4* at 28 dpt. The relative abundances of specific fungal classes were comparable in the roots and rhizosphere of all genotypes (**Fig. 1B**).

**Fig. 1.**
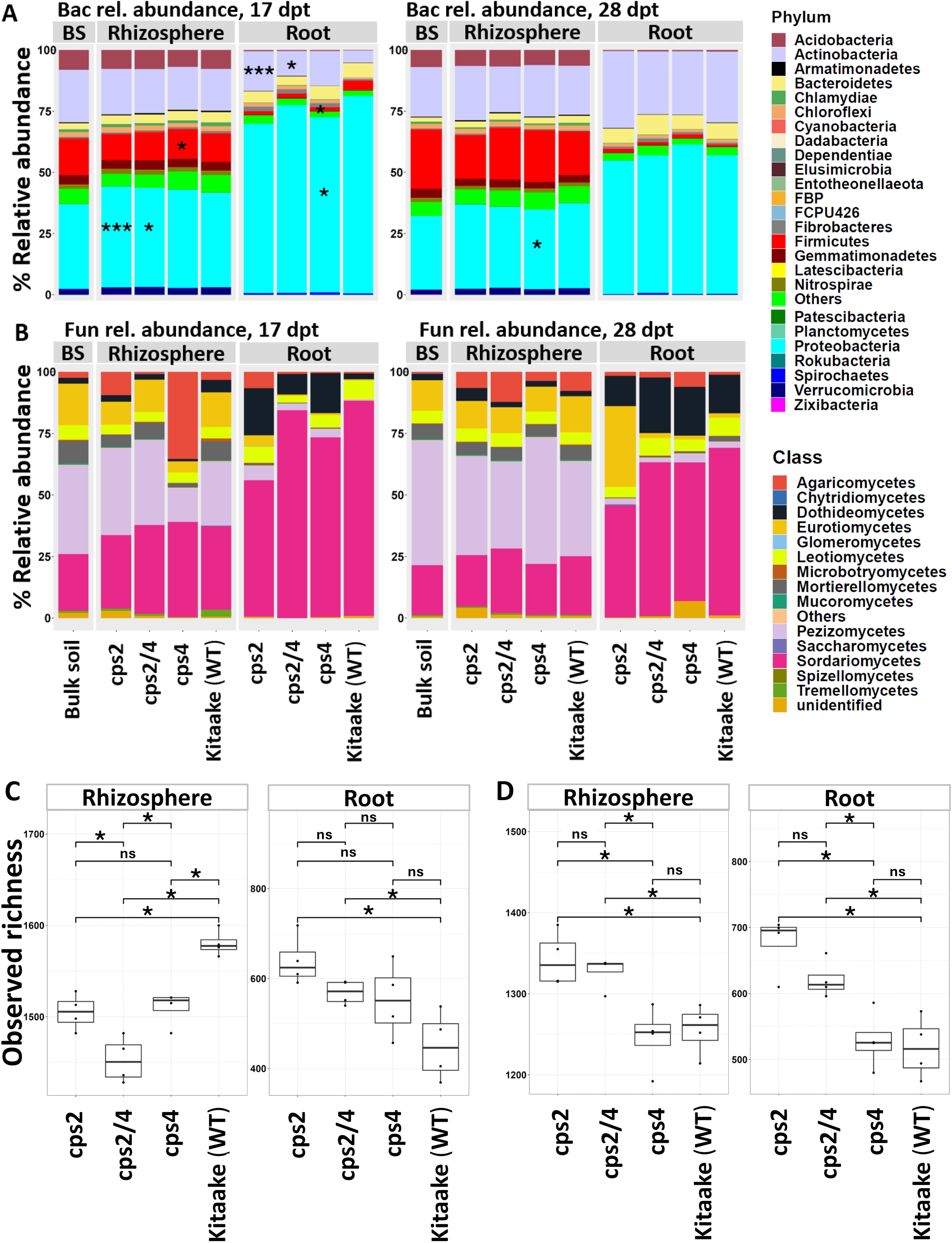
Microbial taxonomic profiles in different rice compartments and growth stages. **A**) Relative abundances of bacterial taxa (phyla-level) in root, rhizosphere and bulk soil at 17 dpt and 28 dpt. **B**) Relative abundances of fungal taxa (class-level) in root, rhizosphere and bulk soil at 17 dpt and 28 dpt. Bacterial observed richness in wildtype and mutant rice lines in the rhizosphere and root compartments at **C**) 17 and **D**) 28 dpt. Microbial taxa with significantly different relative abundances between mutants and wildtype Kitaake were indicated in asterisk in the bar plots (*, P < 0.05; **, P < 0.01, and ***, P < 0.001). Comparisons were performed using Wilcoxon test and P-values were adjusted for multiple comparisons (fdr) using the compare_means() function in the R package ggpubr (v0.6.0).

At the OTU level, bacterial richness in the rhizosphere was lower in all the mutants compared to WT at 17 dpt (**Fig. 1C**), whereas bacterial richness was higher in the *cps2* and *cps2/4* mutants compared to WT in the roots at both sampling times and in the rhizosphere at 28 dpt (**Fig. 1D**). Bacterial richness was lowest in the rhizosphere of *cps2/4* at 17 dpt (**Fig. 1C**) and significantly higher in the rhizosphere and roots of *cps2* than in *cps4* at 28 dpt (**Fig. 1D**). Fungal OTU richness was less responsive to the mutations, but at 17 dpt *cps2* roots had higher fungal richness than WT (**Supplementary Fig. S2**). Moreover, the composition of microbial communities were more distinct in the roots compared with the bulk soil and rhizosphere compartments, with strong effects exerted by both genotype and compartment on bacterial and fungal communities (**Supplementary Fig. S3,4, Table S3)**.

Similarly, at 17 dpt, the root bacterial community composition assessed at OTU level differed significantly between all genotypes. In the rhizosphere, bacterial communities in mutants were all significantly different compared to WT. Also, bacterial communities in the *cps4* mutant were significantly different from *cps2* and *cps2/4* (**Fig. 2A**, **Table 1, Supplementary Fig. S4, Tabl e**).**S** A**4**t 28 dpt, differences between bacterial communities in roots from WT and mutants were not significant (**Fig. 2B**, **Table 1**). Among the mutants, the rhizosphere bacterial communities of *cps2* were significantly different compared with *cps4* (Adonis, R =0.18, P<0.05) and *cps2/4* (Adonis, R =0.18, P<0.05) at 28 dpt (**Supplementary Table S4**). For the fungal communities, the community composition of the WT was significantly different from both of the single mutant lines (*cps2* and *cps4*) in both roots and rhizosphere at 17 dpt and in roots at 28 dpt (**Fig. 2C,D**, **Table 1**). Root fungal communities also were significantly different between *cps2* and *cps2/4* (**Supplementary Table S4**).

**Fig. 2.**
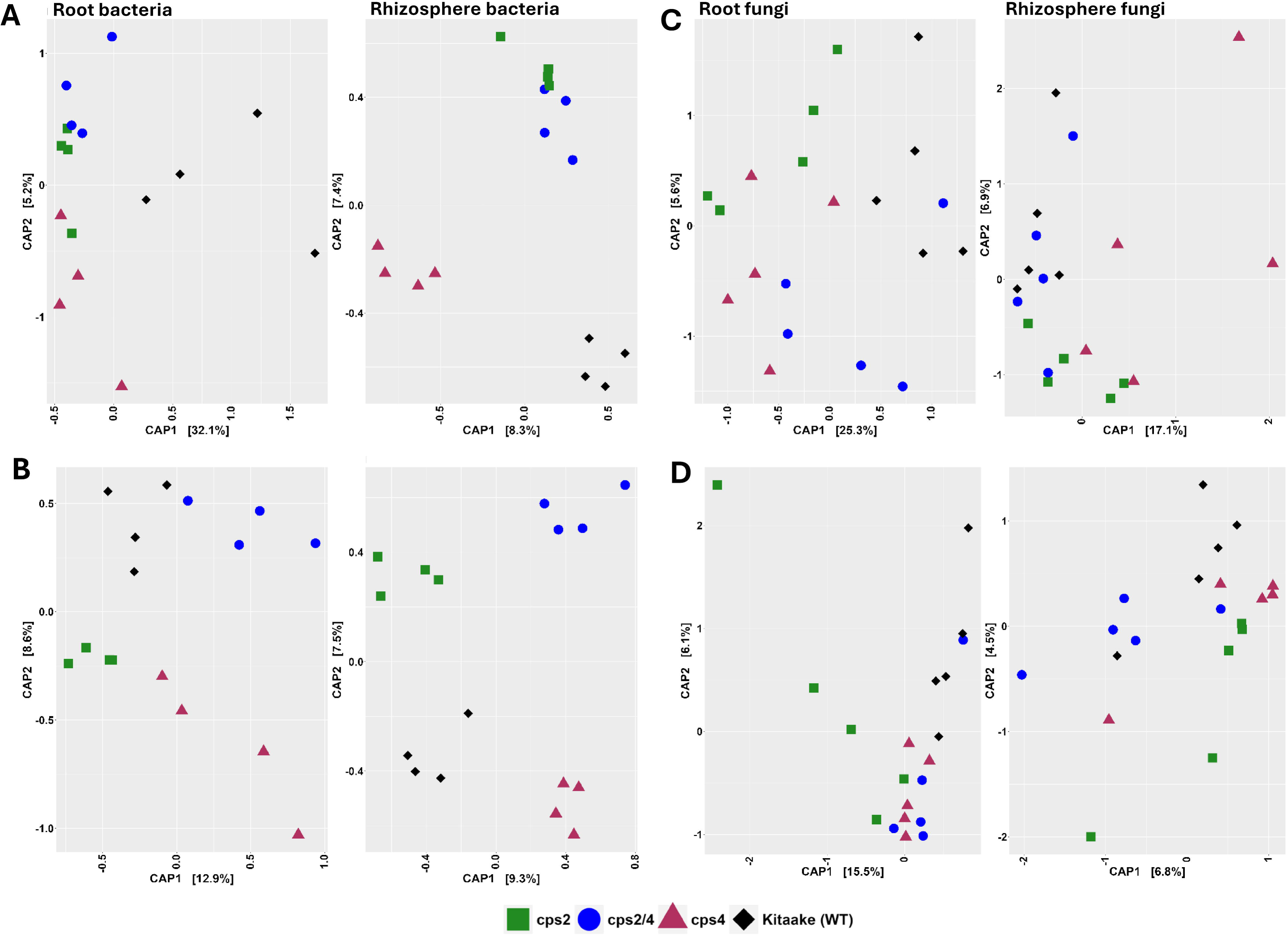
Host genotype effects on microbial communities in the root and rhizosphere compartments. Constrained analysis of principal coordinates (CAP) of bacterial communities in rice root and rhizosphere at **A**) 17 dpt and **B**) 28 dpt and similarly for fungal communities at **C**)17 dpt and **D**) 28 dpt using Bray-Curtis dissimilarity distances.

**Table 1:** PERMANOVA (Pairwise-adonis) between rice mutants and wildtype (Kitaake).

| Community | DPI | Compartment | Factors | R<sup>2</sup> |
| --- | --- | --- | --- | --- |
| Bacteria | 17 | root | cps2_vs_Kitaake | 0.38 * |
|  |  |  | cps4_vs_Kitaake | 0.36* |
|  |  |  | cps2/4_vs_Kitaake | 0.35 * |
|  |  | rhizosphere | cps2_vs_Kitaake | 0.16 ns |
|  |  |  | cps4_vs_Kitaake | 0.18 * |
|  |  |  | cps2/4_vs_Kitaake | 0.15 ns |
|  | 28 | root | cps2_vs_Kitaake | 0.17 ns |
|  |  |  | cps4_vs_Kitaake | 0.19 ns |
|  |  |  | cps2/4_vs_Kitaake | 0.19 ns |
|  |  | rhizosphere | cps2_vs_Kitaake | 0.176 ns |
|  |  |  | cps4_vs_Kitaake | 0.15 ns |
|  |  |  | cps2/4_vs_Kitaake | 0.17 ns |
| Fungal | 17 | root | cps2_vs_Kitaake | 0.38 * |
|  |  |  | cps4_vs_Kitaake | 0.37 ** |
|  |  |  | cps2/4_vs_Kitaake | 0.19 ns |
|  |  | rhizosphere | cps2_vs_Kitaake | 0.29 * |
|  |  |  | cps4_vs_Kitaake | 0.22 * |
|  |  |  | cps2/4_vs_Kitaake | 0.09 ns |
|  | 28 | root | cps2_vs_Kitaake | 0.22 ** |
|  |  |  | cps4_vs_Kitaake | 0.20 * |
|  |  |  | cps2/4_vs_Kitaake | 0.16 ns |
|  |  | rhizosphere | cps2_vs_Kitaake | 0.09 ns |
|  |  |  | cps4_vs_Kitaake | 0.10 ns |
|  |  |  | cps2/4_vs_Kitaake | 0.12 ns |

Significance of test indicated as \*\*, p <0.01; \*, p <0.05. The ns denotes not statistically significant and R<sup>2</sup> is the proportion of variation explained.

### Diterpenoids modulate the relative abundance of several bacterial and fungal OTUs

Next, to identify OTUs that were putatively affected by DPs, we analysed the differences in the abundance of OTUs between the DP mutants and WT using differential abundance analysis. This revealed consistent differences between WT and the three mutant lines; at the individual sampling dates, a number of bacterial and fungal taxa were consistently enriched in WT roots compared to all three mutant lines (**Fig. 3A,B, 4 A**),. **B**At 17 dpt, this applies to e.g. bacterial genera *Kosakonia*, *Paenibacillus*, *Ochrobacter*, *Xanthomonas, Thermomonas*, and *Acidibacter* and fungal genera *Sarcopodium*, and *Fuscoporia* (**Fig. 3A,B**). Also, a few microbial taxa including the bacterial genera *Streptomyces*, *Delftia* and the fungal genera *Clonostachys*, *Waitea* and *Coniochaeta* were only enriched in the single mutants.

**Fig. 3.**
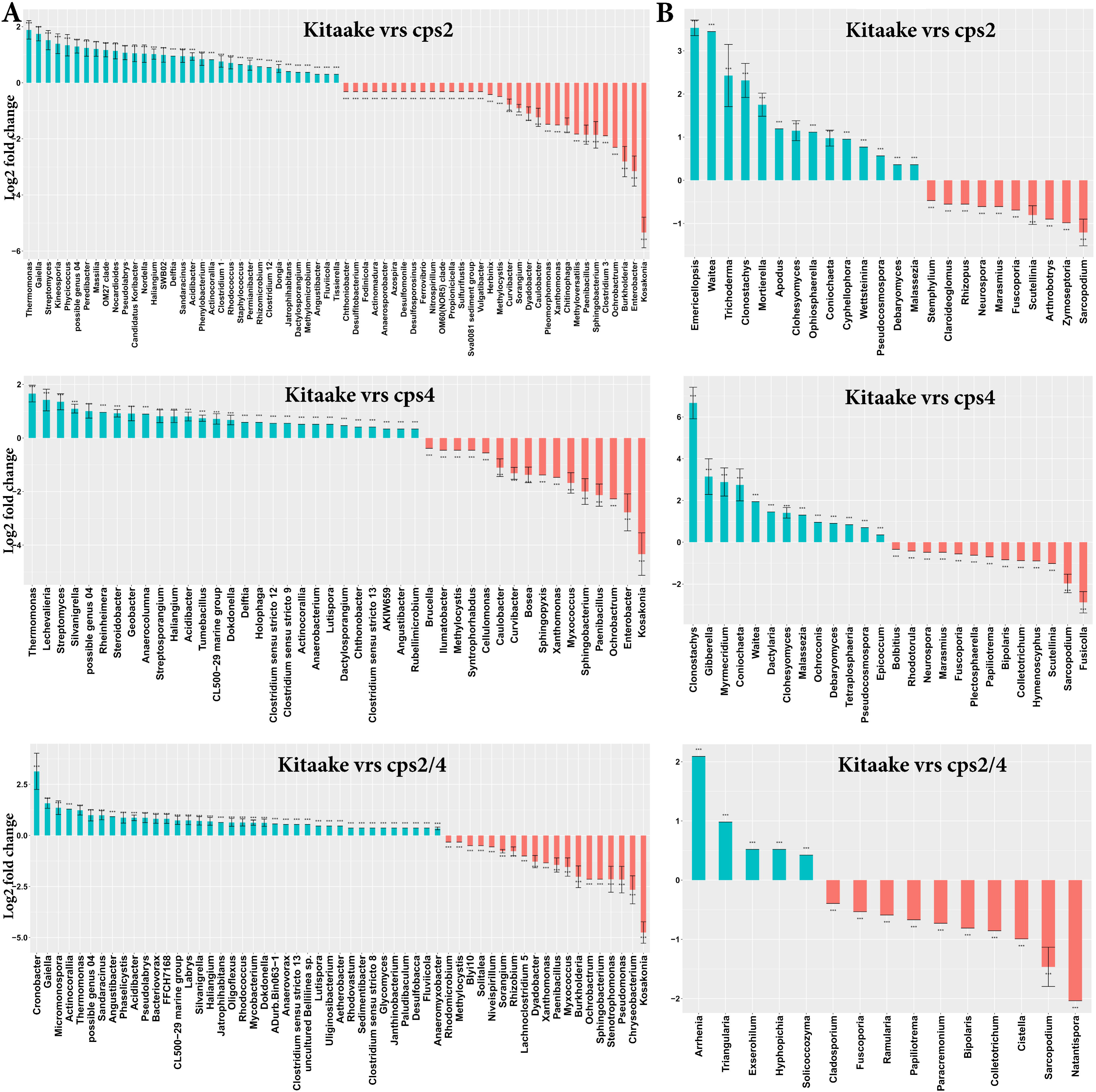
Differentially abundant (log fold change) **A**) bacterial and **B**) fungal genera between roots of rice wildtype and roots of individual mutants at 17 dpt. Data are represented by log fold change (shown as a column), ±SE (shown as error bars) derived from the ANCOM-BC model. ANCOM-BC implements Bonferroni correction to adjust for multiple comparisons. The significance of test is indicated as ***for p < 0.001, ** for p < 0.01 and * for p < 0.05.

Microbial genera that were differentially abundant in WT and mutant roots and rhizospheres were completely different at 17 dpt compared to 28 dpt (**Fig. 3A,B 4A,B, Supplementary Fig. S5** and **S9**). At both time points, the number of enriched bacterial and fungal taxa was higher in the rhizosphere of WT than in the single mutants, suggesting some taxa are attracted to certain DPs (**Supplementary Fig. S5, S6**). At 17 dpt, fungal genera including *Botrytis*, *Wallemia*, *Paraphaeospaeria*, *Stemphylium*, and *Cystofilobasidium* were consistently enriched in the rhizosphere of WT compared with the mutants. *Staphylococcus* was the bacterial genus that was most enriched in roots of all the mutants compared with WT, suggesting that DPs restrict its colonization (**Fig. 4A)**. Fungal taxa *Zymoseptoria*, *Dioszegia* and *Itersonilia* were more abundant in WT than in the single mutants (**Supplementary Fig. S5B**). An overlap of differentially abundant bacterial genera including *Bacteriovorax*, *Pseudoxanthomonas*, *Pedosphaera, Rhodocytophaga* and *Solitalea* were depleted in rhizospheres of *cps2* and *cps4* compared with WT.

**Fig. 4.**
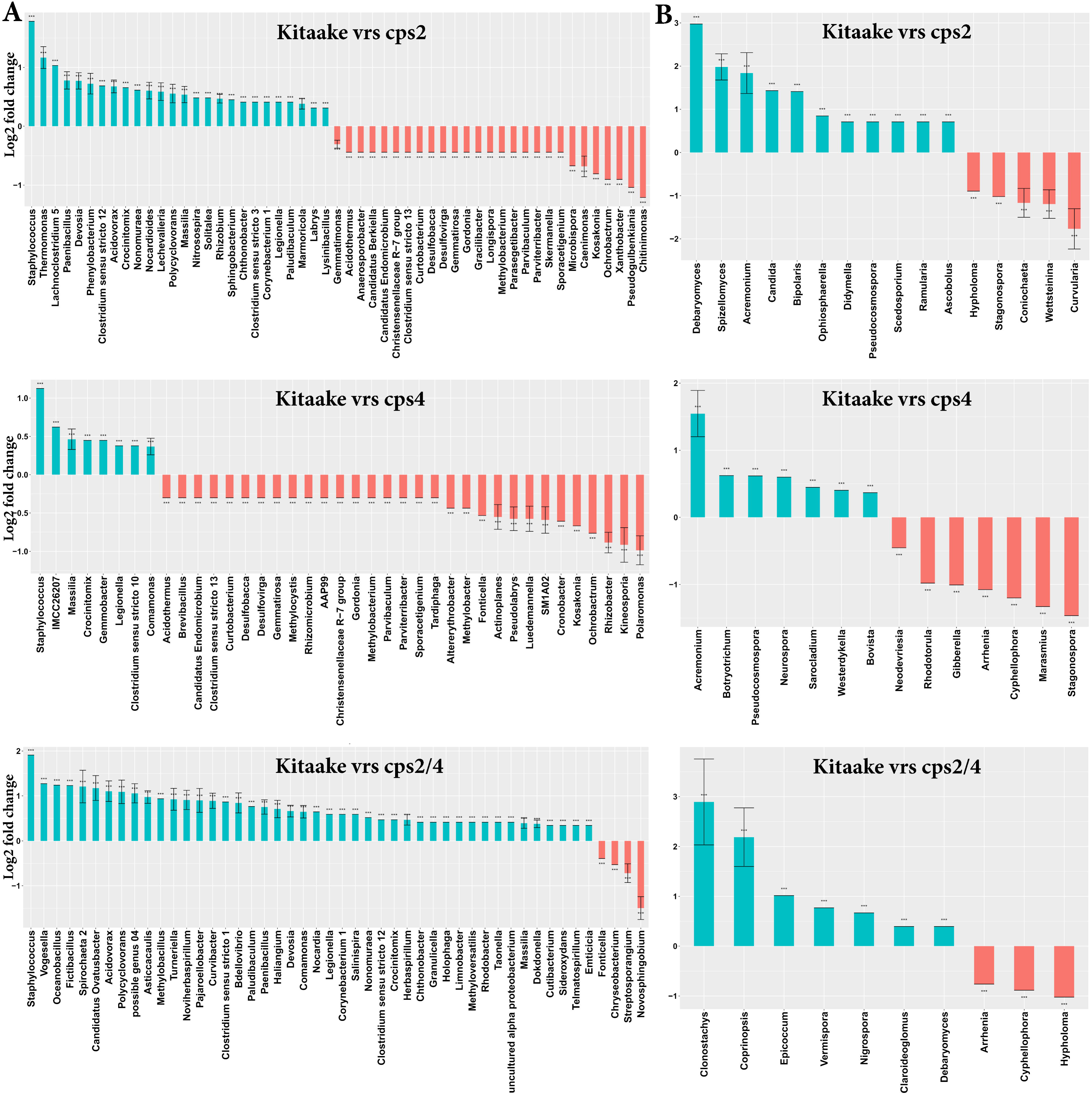
Differentially abundant (log fold change) **A**) bacterial and **B**) fungal genera between roots of rice wildtype and roots of individual mutants at 28 dpt. Data are represented by log fold change (shown as a column), ±SE (shown as error bars) derived from the ANCOM-BC model. ANCOM-BC implements Bonferroni correction to adjust for multiple comparisons. The significance of test is indicated as ***for p < 0.001, ** for p < 0.01 and * for p < 0.05.

Furthermore, differential analysis between the single mutants revealed taxa that were significantly enriched in *cps4* relative to the *cps2* mutant in roots at 17 and 28 dpt and in the rhizosphere at 28 dpt (**Fig. 5, Supplementary Fig. 7**). The differentially abundant taxa were most distinct between root and rhizosphere compartments and between the different time points (**Fig. 5, Supplementary Fig. 7**). For instance, while *Burkholderia*, *Dechloromonas* and *Azospira* were strongly enriched in *cps4*, *Pseudoxanthomonas*, *Chitinomonas* and *Sphingopyxis* were enriched in *cps2*. Also, bacterial taxa including *Burkholderia*, *Dechloromonas* and *Azospira* were highly enriched in roots of *cps4* at 17 dpt, while *Pseudogulbenkiania*, *Azospirillum* and *Chitinimonas* were enriched at 28 dpt. Likewise, a number of differentially abundant fungal taxa were detected when comparing *cps2* and *cps4* mutants (**Fig. 5, Supplementary Fig. 7**).

**Fig. 5.**
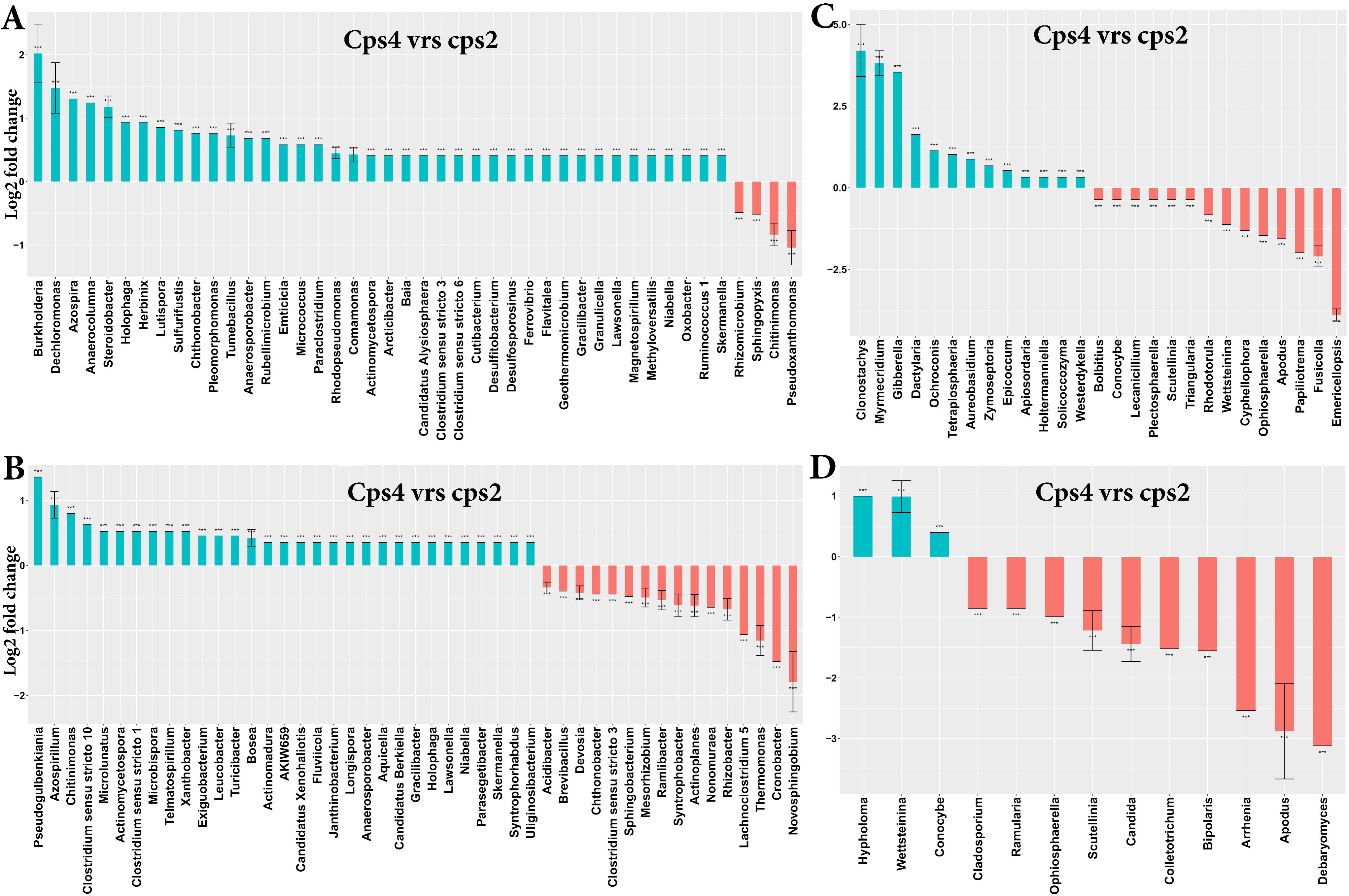
Differentially abundant (log f old change) **A**) bacterial and **B**) fungal genera between roots of rice mutant cps2 and cps4 at 28 dpt. Data are represented by log fold change (shown as a column), ±SE (shown as error bars) derived from the ANCOM-BC model. ANCOM-BC implements Bonferroni correction to adjust for multiple comparisons. The significance of test is indicated as ***for p < 0.001, ** for p < 0.01 and * for p < 0.05.

By comparing each single mutant with the double mutant *cps2/4* we identified several differentially abundant taxa (**Supplementary Fig. S8-11**). Bacterial taxa *Pseudoxanthomonas* and fungal genera *Emericellopsis*, *Papiliotrema*, *Apodus*, *Cyphellophora*, and *Wettsteinina* were consistently enriched in roots of *cps2* compared to *cps4* and *cps2/4* at 17 dpt, and similarly for taxa such as *Novosphingobium*, *Sphingobacterium*, *Brevibacillus* at 28 dpt (**Fig. 5, Supplementary Fig. 8**). *Azospira*, *Lutispora*, *Sulfurifustis*, and *Pleomorphomonas*, were consistently enriched in roots of *cps4* and *cps2/4* compared to *cps2* at 17 dpt (**Fig. 5, Supplementary Fig. 8**).

### Diterpenoids affect associations between microorganisms and nematodes

Next, we explored putative microbial-nematode interactions by performing correlation analysis and also by examining to what extent patterns of correlations varied between the WT and DP mutants. Overall, patterns of correlations between microbial and nematode taxa were very distinct between the rice genotypes, especially between *cps2* and *cps2/4* (**Supplementary Fig. S12**). Only a few bacterial taxa including *Acidobacter*, *Devosia*, *Enterobacter*, *Massilia*, *Stenotrophomonas, Streptomyces* and Burkholderiaceae correlated negatively with the parasitic nematode genus *Meloidogyne* (**Fig. 6A**). Of these, only *Streptomyces* correlated negatively with *Meloidogyne* in all genotypes, whereas the other taxa varied between genotypes. Most positive correlations between bacterial taxa and *Meloidogyne* were not affected by DPs (**Supplementary Fig. S13A**). For instance, the bacterial genera *Sporosarcina*, *Sphingomonas*, *Pseudolabrys*, *Bacillus* and bacterial OTUs assigned to the taxa Methyloligellaceae, Deltaproteobacteria and Acidobacteraceae correlated positively with *Meloidogyne* in most genotypes.

**Fig. 6.**
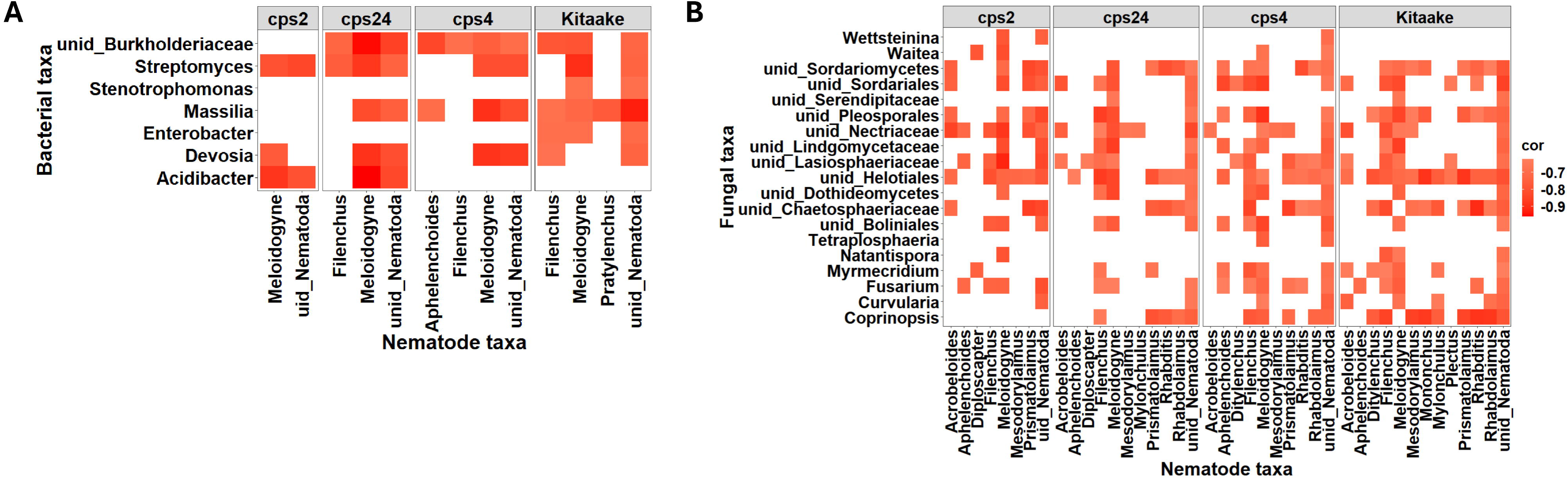
Heat map of microbial-nematode negative associations in rice wildtype and mutants. **A**) Bacterial-nematode negative correlations in rice WT and DP impaired mutants. **B**) Fungal-nematode negative correlations in rice WT and DP impaired mutants. Only microorganisms that negatively correlated with *M. graminicola* were used for the visualization.

A large number of fungal taxa correlated negatively with *Meloidogyne* (**Fig. 6B**). These included *Fusarium*, *Coprinopsis*, *Curvularia*, *Myrmecridium*, *Natantispora*, *Tetraplosphaeria*, *Waitea*, *Wettsteinina* and several OTUs belonging to Sordariomycetes. A higher number of these correlations were observed in WT and *cps4*. In addition, several positive associations were revealed between fungal taxa and *Meloidogyne* in both WT and the mutants (**Supplementary Fig. S3B**).

## Discussion

### Diterpenoids modulate root-associated microbiomes in rice

Previously, we have shown that disruption of rice DP biosynthesis resulted in distinct nematode communities, specifically in the *cps2*, *cps4* and *cps2/4* mutants compared to WT, and most strikingly, the three mutants were more susceptible to *M. graminicola* infection than WT (Desmedt *et al*., 2022). As these DPs are generally known to be antimicrobial, here we ask if DPs also shaped the composition of microbial communities associated with rice roots, and if so, if there are indications that DP-governed changes in nematode communities and rice susceptibility to *M. graminicola* are linked to microbiome changes.

The higher bacterial OTU richness in roots of *cps2* mutants aligns with the higher diversity in microbiomes of *ent*-CDP derived diterpenoid-deficient maize (Murphy *et al*., 2021) and confirms that the antimicrobial activity of these DPs restricts root colonization and/or proliferation of members of the soil microbiota. Although fungal OTU diversity did not vary significantly between DP mutants and WT, disruption of DP synthesis modulated the composition of both bacterial and fungal communities. This corroborates the strong selective effect that exudation of secondary metabolites imposes on root microbiome assembly shown for other bioactive metabolites such as the benzoxazinoids (Kudjordjie *et al*., 2019), flavonoids, glucosinolates and camalexin in other plants (Sikder *et al*., 2022).

We found that the relative abundances of bacterial phyla and fungal classes varied between genotypes and sampling times. At 28 dpt, only *cps4* exhibited any significant difference in rhizosphere communities. At the OTU level, a similar pattern emerged, where the compositional differences between WT and the mutants were more pronounced in the roots than in the rhizosphere and at the earlier (17 dpt) than later (28 dpt) sampling time. These results align with a study which showed that a vast array of triterpenes influenced the assembly of *Arabidopsis* root associated microbiota (Huang *et al*., 2019). Moreover, differences in relative abundances of bacterial taxa in roots between single mutants at 17 dpt, indicate distinct effects of *ent*-CDP derived (*OsCPS2*-dependent) versus *syn*-CDP derived (*OsCPS4*-dependent) DPs in rice microbiome assembly. For instance, the strong enrichment of Actinobacteria in roots of *cps2*, and the higher species richness in both root and rhizosphere of *cps2* than *cps4* suggests that *ent*-CDP derived DPs exert a stronger selection effect on rice bacterial community assembly than *syn*-CDP derived DPs.

As noted, the differences in microbial communities were more evident at the early sampling than at the later sampling. This suggests an early structuring effect of DPs on rice associated microbiomes, which may be caused by high DP secretion in younger plants. Plants are known to produce high quantities of defensive metabolites at early development stages, to ensure protection of seedlings (Kudjordjie *et al*., 2019), and young rice seedlings are known also to exude DPs (Toyomasu *et al*., 2008).

### Diterpenoids affect putative microbe-nematode interactions

Previous work has shown that co-variation in plant production of secondary metabolites and susceptibility to root-knot nematodes may be modulated via microbial responses (Sikder *et al*., 2022). Therefore, we wanted to further explore whether the DP-induced changes in nematode communities, most notably the enrichment of *M. graminicola* in the DP mutants (Desmedt *et al*., 2022), was related to microbial changes between WT and the mutants. Comparing correlations between nematode and microbial taxa across the four rice genotypes, it is apparent that overall patterns of correlations between nematode taxa and individual bacterial and fungal OTUs are distinct. This clearly suggests that DPs play a role in tritrophic inter-kingdom interactions within the root-associated biota. Only a limited number of bacterial taxa correlated with *Meloidogyne*, notably *Streptomyces* which correlated negatively with *Meloidogyne* in all rice genotypes. *Stenotrophomonas* and *Enterobacter* only correlated (negatively) with *Meloidogyne* in WT, and *Massilia* as well as an OTU assigned to Burkholderiaceae correlated negatively with *Meloidogyne* in all genotypes except *cps2*, whereas *Acidibacter* correlated negatively with *Meloidogyne* in the two mutant lines carrying the *cps2* mutation (*cps2* and *cps2/4*). The negative correlations suggest that these bacterial taxa play a role in suppressing *M. graminicola* infectivity and corroborate previous reports on nematode-suppressive activity exerted by strains within these taxa. For instance, *Enterobacter* strains (Duponnois *et al*., 1999; Oliveira *et al*., 2007; Oh *et al*., 2018; Zhao *et al*., 2022), *Stenotrophomonas* strains (Mekete *et al*., 2009; Groover *et al*., 2020), *Burkholderia* species (Zhang *et al*., 2022; Kim *et al*., 2023), and many *Streptomyces* species (Nimnoi and Ruanpanun, 2020; Topalović *et al*., 2020; Hu *et al*., 2022) are known to be nematicidal. In addition, chitinolytic activity of *Massilia* has been suggested to be involved in the suppression of plant parasitic nematodes (Cretoiu *et al*., 2013). In a previous study, we likewise reported negative correlations between *Acidobacter* and *M. incognita* in *Arabidopsis* roots (Sikder *et al*., 2021).

### Diterpenoids affect the abundance of specific bacterial and fungal taxa

Having established that the DP mutants and WT assembled distinct root microbiomes, we aimed to identify bacterial and fungal taxa that were differentially represented in mutant and WT microbiomes. In line with the higher bacterial OTU richness in the mutants, a considerable number of taxa were enriched in the microbiomes of these DP deficient lines, whereas only a limited number of taxa had a higher relative abundance in WT. In general, the comparison between WT and the individual mutants revealed that a few taxa were consistently enriched either in WT or mutants. This consistent pattern suggests that the antimicrobial activity of DPs exert a modulating effect on specific microbial taxa during the establishment of the root microbiome.

Interestingly, specific microbial taxa, for example the bacterial genera *Streptomyces*, *Delftia* and the fungal genera *Clonostachys*, *Waitea* and *Coniochaeta,* were consistently enriched in roots of single mutants but not in the double mutant at 17 dpt. Conversely, a number of microbial taxa were enriched in the roots and rhizosphere of the double mutant at both at 17 dpt and 28 dpt. We speculate that in the double mutant, the disruption of both *OsCPS2* (i.e., DPs derived from *ent*-CDP) and *OsCPS4* (i.e., DPs derived from *syn*-CDP) genes differentially affect individual microbial taxa.

The higher number of differentially abundant bacteria genera identified in roots of *cps4* at 17 dpt and the enrichment of specific fungal taxa in *cps2* roots at 28 dpt, further support the distinct effects of DPs derived from *ent*-CDP and *syn*-CDP branches of the rice DP metabolic network. Lu et al., (Lu *et al*., 2018) reported that *OsCPS2*-dependent DPs were active in rice plant defense against both the fungal pathogen *M. oryzae* and bacterial pathogen *X. oryzae*, while *OsCPS4*-dependent DPs were involved in non-host resistance against the turfgrass fungal pathogen *M. poae*. The DPs dependent on *OsCPS2* are mainly the phytocassanes and abietoryzins, while the *OsCPS4*-dependent DPs include the momilactones and oryzalexin S (Toyomasu *et al*., 2014). The activity of the *ent*-CDP and *syn*-CDP derived DPs against specific fungal species have been widely reported (reviewed in (Valletta *et al*., 2023)). Previous studies reported that momilactones did not inhibit the rice blast fungi *M. oryzae* or *Fusarium fujikuroi*, causative agent of the bakanae ‘foolish seedling’ disease (Xu *et al*., 2012). Also, studies have shown that momilactone B is more active against *Botrytis cinerea*, *Fusarium solani*, *Colletrotrichum gloeosporioides* than momilactone A, whereas neither of these momilactones affect *Fusarium oxysporum* (Fukuta *et al*., 2007). Moreover, the consistent enrichment of specific microbial taxa in the mutants reported here suggests that the depletion of specific DPs enables their establishment in the root (Zarraonaindia *et al*., 2015). Altogether, the present findings demonstrate that DPs have a profound effect on individual microbial taxa, exhibiting some specificity in their interactions, and overall modulating the assembly of rice associated microbiomes.

## Conclusion

Our study provided detailed evidence for various roles of DPs in structuring the root-associated microbiome of rice plants and showed indications of microbiome mediated effects on the parasitic root-knot nematode genus *Meloidogyne*. We found that disruption of the two branches of the rice DP biosynthetic network distinctively affected the assembly of rice root microbiomes. Moreover, DP mutations caused significant shifts in bacterial diversity. Bacterial OTU richness was higher in the *cps2*, and *cps2/4* mutants compared to WT in the roots at both sampling times and in the rhizosphere at 28 dpt. It appears that DPs have a “gatekeeper” role; i.e. DPs block certain microbial taxa from establishment in the rice root-associated microbiome. Interestingly, several bacterial and fungal taxa that correlated negatively with *M. graminicola* or are known to control root-knot nematodes were enriched in WT compared to the mutants. This is interesting, as *M. graminicola* was previously shown to be less abundant in the WT than in the mutant lines. Bacterial taxa that correlated negatively with *M. graminicola*, for example *Streptomyces*, *Enterobacter*, *Xanthomonas*, *Sphingobacterium* and *Paenibacillus*, were strongly enriched in roots of WT relative to the DP mutants at the early growth stages of rice development. These taxa together with the highly enriched genus *Kosakonia* are antagonists of microbial pathogens as well. These findings provide insights into rice-secreted DPs-mediated assembly of root microbiomes and reveal potentially beneficial microbial taxa that seem to be recruited by secreted DPs. Hence these results further support the importance of breeding efforts that target DP synthesis as a promising route towards rice cultivars fit to harnessing pest and pathogen suppressive microbiomes.

## Supplementary figures

**Fig. S1.** (**A**) bacterial and (**B**) fungal sequence reads and OTUs in samples used in this study. Rarefaction curves showing the coverage of (**C**) bacterial and (**D**) fungal OTU richness (species richness in number of OTUs) as a function of sequencing depth (sample size in number of reads).

**Fig. S2.** Bacterial Shannon diversity in root and rhizosphere of WT and mutant rice at **A**) 17 and **B**) 28 dpt. Fungal alpha diversity (Observed and Shannon) in root and rhizosphere of WT and mutant rice at **C**) 17 and **D**) 28 dpt.

**Fig. S3.** Microbial community composition in soil and rice lines. Principal Coordinates Analysis (PCoA) of rice WT and mutants **A**) bacterial and **B**) fungal communities. Analysis was performed using datasets from both 17 and 28 dpt.

**Fig. S4.** Principal Coordinates Analysis of microbial communities in rice WT and mutants using Bray-Curtis dissimilarity distances. PCoA plots of bacterial community in root and rhizosphere at **A**) 17 and **B)** 28 dpt. PCoA plots of fungal community in root and rhizosphere at **A)** 17 and **B)** 28 dpt.

**Fig. S5**. Differentially abundant (log gold change) microbial genera in the rhizosphere of Kitaake (wildtype rice) and mutants. Differentially abundant A) bacterial and B) fungal genera between Kitaake (red) and individual mutants (green) at 17 dpt. Data are represented by log fold change (shown as a column), ±SE (shown as error bars) derived from the ANCOM-BC model. The significance of test is indicated as ***for p < 0.001, ** for p < 0.01 and * for p < 0.05.

**Fig. S6**. Differentially abundant (log gold change) microbial genera in the rhizosphere of Kitaake (wildtype rice) and mutants. Differentially abundant A) bacterial and B) fungal genera between Kitaake (red) and individual mutants (green) at 28 dpt. Data are represented by log fold change (shown as a column), ±SE (shown as error bars) derived from the ANCOM-BC model. The significance of test is indicated as ***for p < 0.001, ** for p < 0.01 and * for p < 0.05.

**Fig. S7.** Differentially abundant (log gold change) microbial genera in the rhizosphere of cps2 (red) and cps4 (green). Differentially abundant A) bacterial and B) fungal genera between Kitaake and individual mutants at 28 dpt. Data are represented by log fold change (shown as a column), ±SE (shown as error bars) derived from the ANCOM-BC model. The significance of test is indicated as ***for p < 0.001, ** for p < 0.01 and * for p < 0.05.

**Fig. S8.** Differentially abundant (log gold change) root bacterial taxa between cps2 (red) and cps2/4 (green) at A)17dpt and B) 28dpt and fungal genera at C) 17dpt and D) 28dpt. Data are represented by log fold change (shown as a column), ±SE (shown as error bars) derived from the ANCOM-BC model. The significance of test is indicated as ***for p < 0.001, ** for p < 0.01 and * for p < 0.05.

**Fig. S9.** Differentially abundant (log gold change) root bacterial taxa between cps4 (red) and cps2/4 (green) at A)17dpt and B) 28dpt and fungal genera at C) 17dpt and D) 28dpt. Data are represented by log fold change (shown as a column), ±SE (shown as error bars) derived from the ANCOM-BC model. The significance of test is indicated as ***for p < 0.001, ** for p < 0.01 and * for p < 0.05.

**Fig. S10.** Differentially abundant (log gold change) rhizosphere bacterial taxa between cps2 (red) and cps2/4 (green) at A)17dpt and B) 28dpt and fungal genera at C) 17dpt and D) 28dpt. Data are represented by log fold change (shown as a column), ±SE (shown as error bars) derived from the ANCOM-BC model. The significance of test is indicated as ***for p < 0.001, ** for p < 0.01 and * for p < 0.05.

**Fig. S11.** Differentially abundant (log gold change) rhizosphere bacterial taxa between cps4 (red) and cps2/4 (green) at A)17dpt and B) 28dpt and fungal genera at C) 17dpt and D) 28dpt. Data are represented by log fold change (shown as a column), ±SE (shown as error bars) derived from the ANCOM-BC model. The significance of test is indicated as ***for p < 0.001, ** for p < 0.01 and * for p < 0.05.

**Fig. S12**. Heat map of microbial-nematode associations in rice WT and mutants. Both microbes that positively and negatively correlated with *M. graminicola* were used for the visualization.

**Fig. S13**. Heat map of microbial-nematode associations in rice WT and mutants. Only microbes that positively correlated with *M. graminicola* were used for the visualization.

## Supplementary Tables

**Table S1:** Primer sets used in this study for the amplification of bacterial 16S rRNA V5-7 genes. Multiplex identifiers (MID) are indicated in bold in forward primer (799F).

**Table S2:** Mean, median and range of reads per compartment for fungal and bacterial libraries at different days post inoculation (DPI).

**Table S3:** Permutation analysis of variance (PERMANOVA) using “adonis” test on Bray-Curtis distance matrices for bacterial and fungal community dissimilarity assessment using 1000 permutations.

**Table S4:** PERMANOVA (pairwise-adonis) between rice mutants cps2 and cps4.

## Acknowledgements

The authors gratefully acknowledge Stefano Sacchi (Plant Health Service, Regione Lombardia, Italy) for collecting field soil for the metabarcoding experiment, and Prof. Bing Yang (University of Missouri) for construction of the *cps* mutant lines.

The work was funded by an Aarhus University Research Foundation Starting Grant (grant no. AUFF-F-2018-7-7) and a grant from the US Department of Agriculture – National Institute of Food and Agriculture (grant no. 2020-67013-32557 to RJP).

## Data availability

The raw bacterial and fungal sequence data has been uploaded to Sequence Read Archive (SRA) bioproject number PRJNA1101406 with accession numbers SUB14384063 and SUB14384196), respectively. The raw nematode sequence data can be assessed using the bioproject number SRA bioproject number PRJNA797929 (Desmedt *et al*., 2022).

## Author contributions

This manuscript was written by ENK, MV and RJP, with substantial input from MN. Nematode infection experiments were performed by WD and metabarcoding analysis was performed by ENK. RJP, TK and WD provided the cps mutants. All authors edited and reviewed the manuscript.

## References

Abarenkov K, Henrik Nilsson R, Larsson K-H, et al. 2010. The UNITE database for molecular identification of fungi--recent updates and future perspectives. The New phytologist 186, 281–5.

Beckers B, Op De Beeck M, Thijs S, Truyens S, Weyens N, Boerjan W, Vangronsveld J. 2016. Performance of 16s rDNA primer pairs in the study of rhizosphere and endosphere bacterial microbiomes in metabarcoding studies. Frontiers in Microbiology 7:650.

Bengtsson-Palme J, Ryberg M, Hartmann M, et al. 2013. Improved software detection and extraction of ITS1 and ITS2 from ribosomal ITS sequences of fungi and other eukaryotes for analysis of environmental sequencing data. Methods in Ecology and Evolution, 914–919.

Bodenhausen N, Horton MW, Bergelson J. 2013. Bacterial Communities Associated with the Leaves and the Roots of Arabidopsis thaliana. PLoS ONE 8(2): e56329.

Caporaso JG, Kuczynski J, Stombaugh J, et al. 2010. QIIME allows analysis of high-throughput community sequencing data. Nature methods 7, 335–6.

Chelius MK, Triplett EW. 2001. The diversity of archaea and bacteria in association with the roots of Zea mays L. Microbial Ecology 41, 252–263.

Chinheya CC, Yobo KS, Laing MD. 2017. Biological control of the rootknot nematode, Meloidogyne javanica (Chitwood) using Bacillus isolates, on soybean. Biological Control 109, 37–41.

Chisholm ST, Coaker G, Day B, Staskawicz BJ. 2006. Host-microbe interactions: Shaping the evolution of the plant immune response. Cell 124, 803–814.

Cretoiu MS, Korthals GW, Visser JHM, Van Elsas JD. 2013. Chitin amendment increases soil suppressiveness toward plant pathogens and modulates the actinobacterial and oxalobacteraceal communities in an experimental agricultural field. Applied and Environmental Microbiology 79, 5291–5301.

Desmedt W, Kudjordjie EN, Chavan SN, et al. 2022. Rice diterpenoid phytoalexins are involved in defence against parasitic nematodes and shape rhizosphere nematode communities. New Phytologist 235(3), 1231–1245.

Dixon RA. 2001. Natural products and plant disease resistance. Nature 411, 843–847.

Duponnois R, Bâ AM, Mateille T. 1999. Beneficial effects of Enterobacter cloacae and Pseudomonas mendocina for biocontrol of Meloidogyne incognita with the endospore-forming bacterium Pasteuria penetrans. Nematology 1, 95–101.

Fukuta M, Xuan TD, Deba F, Tawata S, Khanh TD, Chung IM. 2007. Comparative efficacies in vitro of antibacterial, fungicidal, antioxidant, and herbicidal activities of momilatones A and B. Journal of Plant Interactions 2, 245–251.

Groover W, Held D, Lawrence K, Carson K. 2020. Plant growth-promoting rhizobacteria: a novel management strategy for Meloidogyne incognita on turfgrass. Pest Management Science 76, 3127–3138.

Hu Q, Yang M, Bo T, Li Y, Wu C, Mo M, Liu Y. 2022. Soluble macromolecules from two Streptomyces strains with potent nematicidal activity against Meloidogyne incognita. Rhizosphere 22, 100529.

Huang AC, Jiang T, Liu YX, Bai YC, Reed J, Qu B, Goossens A, Nützmann HW, Bai Y, Osbourn A. 2019. A specialized metabolic network selectively modulates Arabidopsis root microbiota. Science 364, eaau6389.

Ihrmark K, Bödeker ITM, Cruz-Martinez K, et al. 2012. New primers to amplify the fungal ITS2 region--evaluation by 454-sequencing of artificial and natural communities. FEMS microbiology ecology 82, 666–77.

Iqbal M, Dubey M, Mcewan K, Menzel U, Franko MA, Viketoft M, Jensen DF, Karlsson M. 2018. Evaluation of clonostachys rosea for control of plant-parasitic nematodes in soil and in roots of carrot and wheat. Phytopathology 108, 52–59.

Jeandet P, Courot E, Clément C, Ricord S, Crouzet J, Aziz A, Cordelier S. 2017. Molecular Engineering of Phytoalexins in Plants: Benefits and Limitations for Food and Agriculture. Journal of Agricultural and Food Chemistry 65, 2643–2644.

Kariya K, Fujita A, Ueno M, Yoshikawa T, Teraishi M, Taniguchi Y, Ueno K, Ishihara A. 2023. Natural variation of diterpenoid phytoalexins in rice: Aromatic diterpenoid phytoalexins in specific cultivars. Phytochemistry 211, 113708.

Kariya K, Ube N, Ueno M, Teraishi M, Okumoto Y, Mori N, Ueno K, Ishihara A. 2020. Natural variation of diterpenoid phytoalexins in cultivated and wild rice species. Phytochemistry 180, 112518.

Kaur S, Samota MK, Choudhary M, Choudhary M, Pandey AK, Sharma A, Thakur J. 2022. How do plants defend themselves against pathogens-Biochemical mechanisms and genetic interventions. Physiology and Molecular Biology of Plants 28, 485–504.

Kim JH, Lee BM, Kang MK, Park DJ, Choi IS, Park HY, Lim CH, Son KH. 2023. Assessment of nematicidal and plant growth-promoting effects of Burkholderia sp. JB-2 in root-knot nematode-infested soil. Frontiers in Plant Science 14, 1–13.

Kudjordjie EN, Sapkota R, Steffensen SK, Fomsgaard IS, Nicolaisen M. 2019. Maize synthesized benzoxazinoids affect the host associated microbiome. Microbiome, 1–17.

Li X, Chou MY, Bonito GM, Last RL. 2023. Anti-fungal bioactive terpenoids in the bioenergy crop switchgrass (Panicum virgatum) may contribute to ecotype-specific microbiome composition. Communications Biology 6, 1–12.

Lin H, Peddada S Das. 2020. Analysis of compositions of microbiomes with bias correction. Nature Communications 11, 1–11.

Lu X, Zhang J, Brown B, et al. 2018. Inferring roles in defense from metabolic allocation of rice diterpenoids. Plant Cell 30, 1119–1131.

Martin M. 2011. Cutadapt removes adapter sequences from high-throughput sequencing reads. EMBnet.journal 17, 10.

McMurdie PJ, Holmes S, Kindt R, Legendre P, O’Hara R. 2013.phyloseq: An R Package for Reproducible Interactive Analysis and Graphics of Microbiome Census Data. (M Watson, Ed.). PLoS ONE 8, e61217.

Mekete T, Hallmann J, Kiewnick S, Sikora R. 2009. Endophytic bacteria from ethiopian coffee plants and their potential to antagonise meloidogyne incognita. Nematology 11, 117– 127.

Murphy KM, Edwards J, Louie KB, Bowen BP, Sundaresan V, Northen TR, Zerbe P. 2021. Bioactive diterpenoids impact the composition of the root-associated microbiome in maize (Zea mays). Scientific Reports 11, 1–13.

Murphy KM, Zerbe P. 2020. Specialized diterpenoid metabolism in monocot crops: Biosynthesis and chemical diversity. Phytochemistry 172, 112289.

Nimnoi P, Ruanpanun P. 2020. Suppression of root-knot nematode and plant growth promotion of chili (Capsicum flutescens L.) using co-inoculation of Streptomyces spp. Biological Control 145, 104244.

Oh M, Han JW, Lee C, Choi GJ, Kim H. 2018. Nematicidal and plant growth-promoting activity of enterobacter asburiae HK169: Genome analysis provides insight into its biological activities. Journal of Microbiology and Biotechnology 28, 968–975.

Oksanen AJ, Blanchet FG, Friendly M, et al. 2020. Package ‘vegan’. https://cran.r-project.org, https://github.com/vegandevs/vegan

Oliveira DF, Campos VP, Amaral DR, Nunes AS, Pantaleão JA, Costa DA. 2007. Selection of rhizobacteria able to produce metabolites active against Meloidogyne exigua. European Journal of Plant Pathology 119, 477–479.

Otomo K, Kenmoku H, Oikawa H, König WA, Toshima H, Mitsuhashi W, Yamane H, Sassa T, Toyomasu T. 2004. Biological functions of ent-and syn-copalyl diphosphate synthases in rice: Key enzymes for the branch point of gibberellin and phytoalexin biosynthesis. Plant Journal 39, 886–893.

Poveda J, Abril-Urias P, Escobar C. 2020. Biological Control of Plant-Parasitic Nematodes by Filamentous Fungi Inducers of Resistance: Trichoderma, Mycorrhizal and Endophytic Fungi. Frontiers in Microbiology 11, 1–14.

Prisic S, Xu M, Wilderman PR, Peters RJ. 2004. Rice contains two disparate ent-copalyl diphosphate synthases with distinct metabolic functions. Plant physiology 136, 4228–4236.

Quast C, Pruesse E, Yilmaz P, Gerken J, Schweer T, Yarza P, Peplies J, Glöckner FO. 2013. The SILVA ribosomal RNA gene database project: Improved data processing and web-based tools. Nucleic Acids Research 41, 590–596.

R Core Team. 2022.R⍰: A Language and Environment for Statistical Computing.

Robinson MD, McCarthy DJ, Smyth GK. 2010. edgeR: A Bioconductor package for differential expression analysis of digital gene expression data. Bioinformatics 26, 139–140.

Rognes T, Flouri T, Nichols B, Quince C, Mahé F. 2016. VSEARCH: a versatile open source tool for metagenomics. PeerJ 4, e2584.

Sakamoto T, Miyura K, Itoh H, et al. 2004. Erratum: An overview of gibberellin metabolism enzyme genes and their related mutants in rice (Plant Physiology (2004) 134 (1642-1653)). Plant Physiology 135, 1863.

Schmelz EA, Huffaker A, Sims JW, Christensen SA, Lu X, Okada K, Peters RJ. 2014. Biosynthesis, elicitation and roles of monocot terpenoid phytoalexins. Plant Journal 79, 659– 678.

Sikder MM, Vestergård M, Kyndt T, Fomsgaard IS, Kudjordjie EN, Nicolaisen M. 2021. Benzoxazinoids selectively affect maize root-associated nematode taxa. Journal of Experimental Botany 72, 3835–3845.

Sikder MM, Vestergård M, Kyndt T, Topalović O, Kudjordjie EN, Nicolaisen M. 2022. Genetic disruption of Arabidopsis secondary metabolite synthesis leads to microbiome-mediated modulation of nematode invasion. ISME Journal, 1–12.

Topalović O, Hussain M, Heuer H. 2020. Plants and Associated Soil Microbiota Cooperatively Suppress Plant-Parasitic Nematodes. Frontiers in Microbiology 11, 1–15.

Toyomasu T, Kagahara T, Okada K, Koga J, Hasegawa M, Mitsuhashi W, Sassa T, Yamane H. 2008. Diterpene phytoalexins are biosynthesized in and exuded from the roots of rice seedlings. Bioscience, Biotechnology and Biochemistry 72, 562–567.

Toyomasu T, Usui M, Sugawara C, et al. 2014. Reverse-genetic approach to verify physiological roles of rice phytoalexins: Characterization of a knockdown mutant of OsCPS4 phytoalexin biosynthetic gene in rice. Physiologia Plantarum 150, 55–62.

Valletta A, Iozia LM, Fattorini L, Leonelli F. 2023. Rice Phytoalexins: Half a Century of Amazing Discoveries; Part I: Distribution, Biosynthesis, Chemical Synthesis, and Biological Activities. Plants 12, 260.

VanEtten HD, Mansfield JW, Bailey JA, Farmer EE. 1994. Two Classes of Plant Antibiotics: Phytoalexins versus ‘Phytoanticipins’. The Plant Cell 6, 1191.

Wickham H. 2009. ggplot2: Elegant Graphics for Data Analysis. Springer-Verlag New York.

Xu M, Galhano R, Wiemann P, et al. 2012. Genetic evidence for natural product-mediated plant-plant allelopathy in rice (Oryza sativa). New Phytologist 193, 570–575.

Xu M, Hillwig ML, Prisic S, Coates RM, Peters RJ. 2004. Functional identification of rice syn-copalyl diphosphate synthase and its role in initiating biosynthesis of diterpenoid phytoalexin/allelopathic natural products. Plant Journal 39, 309–318.

Yamane H. 2013. Biosynthesis of phytoalexins and regulatory mechanisms of it in rice. Bioscience, Biotechnology and Biochemistry 77, 1141–1148.

Zarraonaindia I, Owens SM, Weisenhorn P, et al. 2015. The soil microbiome influences grapevine-associated microbiota. mBio 6, 1–10.

Zhang J, Li R, Xu M, Hoffmann RI, Zhang Y, Liu B, Zhang M, Yang B, Li Z, Peters RJ. 2021. A (conditional) role for labdane-related diterpenoid natural products in rice stomatal closure. New Phytologist 230, 698–709.

Zhang R, Ouyang J, Xu X, et al. 2022. Nematicidal Activity of Burkholderia arboris J211 Against Meloidogyne incognita on Tobacco. Frontiers in Microbiology 13, 1–10.

Zhao Y, Yuan Z, Wang S, et al. 2022. Gene sdaB Is Involved in the Nematocidal Activity of Enterobacter ludwigii AA4 Against the Pine Wood Nematode Bursaphelenchus xylophilus. Frontiers in Microbiology 13, 1–18.

